# Hysteresis in the selective synchronization of brain activity to musical rhythm

**DOI:** 10.1101/696914

**Authors:** Tomas Lenc, Peter E. Keller, Manuel Varlet, Sylvie Nozaradan

## Abstract

When listening to musical rhythm, people tend to spontaneously perceive and move along with a periodic pulse-like meter. Moreover, perception and entrainment to the meter show remarkable stability in the face of dynamically changing rhythmic structure of music, even when acoustic cues to meter frequencies are degraded in the rhythmic input. Here we show that this perceptual phenomenon is supported by a selective synchronization of endogenous brain activity to the perceived meter, and that this neural synchronization is significantly shaped by recent context, especially when the incoming input becomes increasingly ambiguous. We recorded the EEG while non-musician and musician participants listened to nonrepeating rhythmic sequences where acoustic cues to meter frequencies either gradually decreased (from regular to ambiguous) or increased (from ambiguous to regular). We observed that neural activity selectively synchronized to the perceived meter persisted longer when the sequence gradually changed from regular to ambiguous compared to the opposite, thus demonstrating hysteresis in the neural processing of a dynamically changing rhythmic stimulus. This dependence on recent context was weaker in the neural responses of musicians, who also showed greater ability to tap along with a regular meter irrespective of stimulus ambiguity, thus reflecting greater stability relative to current and recent stimulus in musicians. Together, these asymmetric context effects demonstrate how the relative contribution of incoming and prior signals is continuously weighted to shape neural selection of functionally-relevant features and guide perceptual organization of dynamic input.

**Significance statement:** When listening to musical rhythm, people tend to spontaneously perceive and move along with a periodic pulse-like meter. Moreover, perception and entrainment to the meter seem to show remarkable stability in the face of dynamically changing rhythmic structure of music. Here we show that this is supported by a selective synchronization of brain activity at meter frequencies. This selective neural synchronization persists longer when a nonrepeating sequence gradually transforms from a regular to an ambiguous rhythm compared to the opposite. This asymmetric context effect suggests that the brain processes rhythm based on a flexible combination of sensory and endogenous information. Such continuously updated neural emphasis on meter periodicities might therefore guide robust perceptual organization of a dynamic rhythmic input.

## Introduction

One of the biggest challenges in understanding brain function is to explain how coherent representations of the external world are built and maintained based on continuously changing and ambiguous sensory input. Such flexibility and stability is thought to result from a dynamic process whereby the relative contribution of sensory input and endogenous signals is continuously updated (de Lange et al., 2018). This dynamic phenomenon is well illustrated with musical meter, which is often considered a cornerstone of temporal prediction and sensory-motor synchronization with music (Toiviainen et al., 2010; Vuust et al., 2018). Meter perception usually refers to the ability to spontaneously perceive and dance along a nested structure of periodic pulse-like beats when listening to music (London, 2004). Importantly, meter perception arises with a wide range of rhythmic inputs, from strictly metronomic to highly irregular rhythms, thus showing remarkable flexibility and invariance in the mapping between perceptual template and incoming acoustic structure (Repp et al., 2008; Witek et al., 2014; Large et al., 2015).

A growing body of evidence suggests that meter processing is related to a selective synchronization of neural activity at meter periodicities (Nozaradan et al., 2017a), the term synchronization referring here to temporal coordination or phase-locking between two signals, namely activity of large pools of neurons and periodic meter (Lenc et al., 2019). This preferential synchronization of brain activity therefore leads to selective enhancement of frequencies corresponding to the perceived meter, relative to other frequencies that are unrelated to the perceived meter but can be nonetheless prominent in the acoustic input (Nozaradan et al., 2011, 2012; Tal et al., 2017). This transformation of the rhythmic stimulus has been observed within the human auditory cortex (Nozaradan et al., 2016a, 2018), and possibly involves functional connections within an extended cortico-subcortico-cortical network supporting the processing of rhythmic input (Nozaradan et al., 2017b). However, how sensory and endogenous signals are continuously weighted to build this neural representation of rhythm remains unknown. The current study addresses this question by directly testing the influence of recent history of auditory stimulation on the selective neural synchronization to the perceived meter.

Stimulus history, that is, recent context, plays a key role in supporting stable perception in the face of ambiguous sensory input (Snyder et al., 2015). Particularly important for perceptual stability are attractive context effects which bias perception of the current sensory input towards recently encountered stimuli. Such effects have been reported in the auditory (Arzounian et al., 2017; Chambers et al., 2017) and visual modality (Cicchini et al., 2014, 2017; Fischer and Whitney, 2014; Manassi et al., 2017; Suárez-Pinilla et al., 2018), and have also been observed for the reproduction of single temporal intervals (Cicchini et al., 2012). Similar mechanisms are arguably at play during meter perception (London, 2004). Specifically, once a stable metric representation has been established, it tends to withstand ambiguities produced by the continuously changing rhythmic surface of music (Cooper and Meyer, 1963; Lerdahl and Jackendoff, 1983). Moreover, persistence of the metric percept induced by a recent stimulus is compatible with models suggesting that meter representation emerges through coupling between the rhythmic input and a network of non-linear self-sustaining oscillators (Large and Palmer, 2002).

Here, to test the impact of recent context on meter processing, we created auditory sequences gradually changing from regular (onset structure providing prominent cues to a given meter) to syncopated rhythm (irregular onset structure completely ambiguous with respect to the given meter). We also created flipped versions of these sequences, yielding sequences gradually changing from syncopated to regular. EEG activity was recorded from participants while listening to these sequences. After the EEG session, participants were asked to tap along with the perceived pulse of an additional set of sequences constructed with the same algorithm as those used in the EEG session. This behavioral measure therefore indicated the induced metric periodicities across both sets of sequences. Because the envelope spectra of the stimuli were strictly identical across the original and flipped sequences, identical EEG spectra across both sequence directions would suggest the absence of an effect of context. Alternatively, different EEG spectra across the two sequence directions would provide direct evidence for context-dependent neural representation of rhythm. This context effect would be informative about how the relative contribution of sensory and endogenous signals continuously shape neural selection of functionally-relevant features to guide perceptual organization of dynamic input. We compared groups of musicians and non-musicians, with the hypothesis that formal musical training, by providing strong preexisting perceptual templates of meter, would decrease sensitivity to ambiguity of the rhythmic input and thus also to recent context.

## Methods

### Participants

Thirty-two healthy volunteers participated in the study after providing written informed consent. The sample consisted of a group of individuals with no formal musical training (N = 16, mean age = 21.1 y, SD = 5.1 y, 9 females), and a group of musically trained participants (N = 16, mean age = 24.1 y, SD = 5.4 y, 13 females) with various levels of musical training (mean = 7.2 y, SD = 4.9 y). All participants reported normal hearing and no history of neurological or psychiatric disease. The study was approved by the Research Ethics Committee of Western Sydney University.

### Auditory stimulation

We created all possible rhythmic patterns consisting of twelve 200-ms events, where 8 events were sounds (440 Hz pure tone, 10 ms linear onset and offset ramp) and 4 events were silences. After removing phase-shifted versions of the same pattern, this resulted in 43 unique patterns. To quantify how well the arrangement of sound events matched a regular, periodic meter, each pattern was analyzed with a model of syncopation proposed by Longuet-Higgins and Lee (1984), as implemented in the synpy package (Song et al., 2015). The syncopation scores were calculated assuming metrical structure with nested grouping of the individual events by 2, 2, and 3 (i.e. pulses with rates corresponding to 2, 4, and 12 events respectively, such as in a 3/4 meter). Given these particular pulse rates (i.e. meter frequencies), there were 12 possible ways to align the metric template with each analyzed rhythmic pattern (i.e. 12 meter phases). For patterns with highly regular arrangement of sound intervals, the close match of the rhythmic structure and periodic meter for certain alignments would necessarily result in poor match for other alignments. On the other hand, for patterns with more ambiguous structure, there would be no single alignment resulting in close match between the rhythmic structure and the metric template. Therefore, the regularity of each rhythmic pattern was assessed by calculating the range of syncopation scores across the 12 possible meter phases (the highest minus the lowest score). This value also describes the degree of phase-stability of the meter induced by each pattern. While patterns with large range of syncopation strongly encourage perception of particular meter phases over others, there is no such preference for patterns with small syncopation ranges (Povel and Essens, 1985; Fitch and Rosenfeld, 2007). Based on this analysis, the 43 patterns were then categorized into 8 groups (syncopation ranges {8,7,6,5,4,3,2,1}, omitting the single rhythm with range of 9), i.e., from high syncopation range (regular patterns) to low syncopation range (ambiguous patterns).

Next, we created 57.6-sec long sequences, by concatenating 24 patterns randomly chosen (with repetition) from the 43 patterns in such a way that the range of syncopation decreased continuously throughout the sequence. To do so, three different patterns were chosen in each syncopation range group from 8 to 1, thus yielding 3 x 8 = 24 patterns per sequence in total, with gradually decreasing meter stability. After randomly choosing a pattern within the desired syncopation-range group, its particular phase was chosen so that the syncopation score continuously increased throughout the sequence, i.e. increasing complexity with respect to the meter induced by the patterns (syncopation scores {−1,−1,0,1,2,3,4,4} for the eight syncopation range groups). This resulted in a sequence that gradually transformed from regular to syncopated without structural changes likely to trigger mental phase-shifts that would markedly change the perceived syncopation from high to low (e.g. Fitch and Rosenfeld, 2007).

In order to construct sequences with a gradual change in the other direction (from syncopated to regular), we created a time-inverted version of each 57.6-s sequence, so that the first event became the last event. However, simply flipping a rhythmic pattern in time can yield different syncopation score for the original and inverted version because the on-beat positions in the original pattern become the off-beat positions in the inverted pattern. Moreover, listeners have a tendency to perceive the first sound event of a rhythmic pattern as a strong beat (Parncutt, 1994), which could bias their perception of the inverted sequences towards a meter that is misaligned with the accent structure of the stimulus. To remedy this, we added a train of 12 sound events to the start of each sequence before performing the inversion. We also removed the last 3 events from the sequence and added a train of 15 sound events at the end of the sequence (see Figure 1). Syncopation scores were then calculated by taking 24×12 events starting at event 13 for both the original sequence and the inverted sequence. These longer trains of sound events were also added to yield similar levels of neural adaptation at the start of the original and inverted sequences (Fruhstorfer et al., 1970; Budd et al., 1998). This prevented spurious differences in the neural response between sequence directions, which could otherwise arise due to increased transient responses to sound events at the beginning of each sequence.

**Figure 1.**
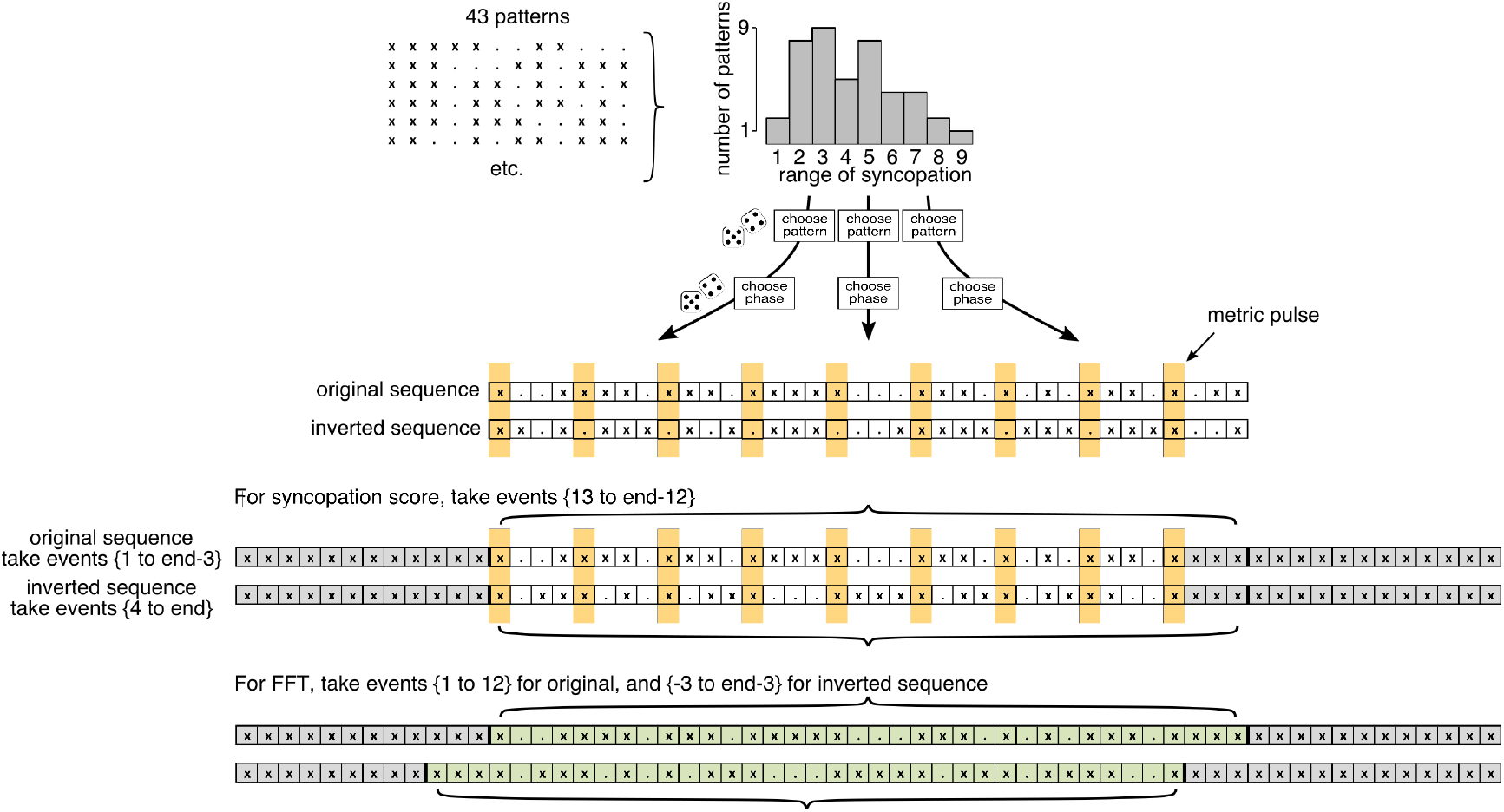
Illustration of the sequence generation method. (Top) Examples of individual constituent patterns used to construct the sequences. Each pattern contains 8 sounds (depicted as “x”), and 4 silences (depicted as “.”). The patterns were categorized based on the range of syncopation across all 12 possible meter phases (calculated separately for each pattern). Sequences were constructed by randomly sampling patterns according to their range of syncopation. After a pattern was selected, its particular phase (i.e. starting point) was sampled according to the particular syncopation score required. (Middle) Example of a regular sequence consisting of 3 patterns (for illustration purposes). Yellow stripes mark pulse positions. Inverting the sequence in time moves the sounds away from the pulse positions. (Bottom) To remedy this, the last 3 events of the original sequence (i.e. the first 3 events of the inverted sequence) are discarded, and the sequence is padded with sound events. Curly brackets mark the parts taken for syncopation score calculation, and for analysis with the cochlear model.

Fifteen unique sequences and their respective inverted versions were generated, forming stimuli for two experimental conditions: the original sequences that evolved from low to high syncopation (regular-to-syncopated condition) and their inverted versions that progressed from high to low syncopation (syncopated-to-regular condition). Five additional sequences and their inverted versions were constructed for the tapping session. The auditory stimuli were created in Matlab R2016b (The MathWorks, Natick, MA) and presented binaurally through insert earphones (ER-2; Etymotic Research, Elk Grove Village, IL) at 75 dB SPL using PsychToolbox, version 3.0.14 (Brainard, 1997).

### Stimulus analysis

#### Syncopation score

To calculate the evolution of syncopation scores across the generated sequences, the sequences were divided into 14.4-sec-long segments (72 events per segment) with 50% overlap, yielding 7 distinct segments per sequence. To evaluate whether the corresponding segments in the original and inverted sequences differed in their degree of syncopation, syncopation scores proposed by Longuet-Higgins and Lee (1984) were calculated for each segment, assuming nested grouping by 2×2 events (i.e., embedded pulses at the rates of 2 and 4 events). This corresponded to the meter used during sequence construction without the slowest pulse, as the individual constituent patterns were not repetitively looped in the sequence. Syncopation scores were compared across conditions using a linear mixed model with Direction (regular-to-syncopated vs. syncopated-to-regular) and Segment (1-7) as fixed effects. In this test and further statistical tests, for all models including the factor Segment as a fixed effect, the order of segments from the syncopated-to-regular condition was always reversed in order to compare responses to the exact inverted versions of the same rhythmic stimulus.

The analysis of the syncopation scores calculated for the 15 stimulus sequences used in the EEG session yielded a significant interaction between the factor Direction and Segment (F = 4.7, P = 0.0002, BF_10_ > 100), suggesting that across trials, inversion of the sequences affected only certain segments. Post-hoc contrasts revealed that the syncopation score was significantly higher for the regular-to-syncopated condition in segment 4 (β = 2.2, t = 2.97, P = 0.02) and 5 (β = 2.4, t = 3.34, P = 0.01). Even though these results suggest that the inversion procedure did not perfectly preserve the theoretically expected amount of syncopation in the sequences, the direction of the effect was opposite to the effect of context we expected to find in the EEG responses. In other words, according to the syncopation scores, there should be more salient cues to the meter in the middle segments in the syncopated-to-regular condition.

The procedure used to construct the auditory stimuli in the current study was based on variations in syncopation that assumed a specific metrical interpretation (nested grouping by 2×2 events). However, there are other possible metrical interpretations of the sequences, which were not considered during stimulus construction. To ensure that the stimulus sequences did indeed change, in theory, from an unambiguous 2×2 meter into highly syncopated sequences instead of converging onto a different meter, we calculated the evolution of syncopation scores across the sequence for two other possible metrical interpretations (nested grouping by 2×3, corresponding to pulse rates of 3 and 6 events; and by 3×2, corresponding to pulse rates of 2 and 6 events). If the sequences modulated into a different meter, then we would expect to find monotonically decreasing syncopation scores for that meter as the sequence progressed from regular to syncopated. As shown in Figure 2, this was not the case for the two other tested meters, further justifying the stimulus construction method that was used.

**Figure 2.**
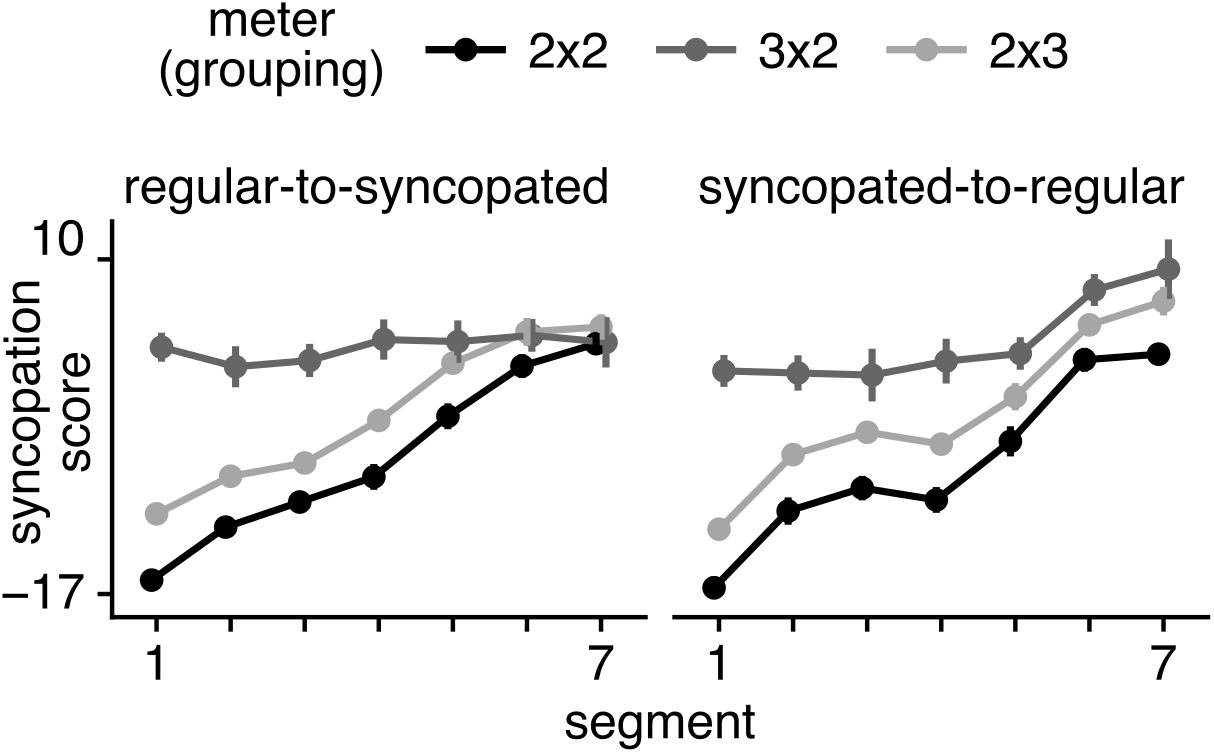
Evolution of syncopation scores across segments, assuming different metrical interpretations of the sequences (grouping by 2×2, 3×2, and 2×3 events). Syncopation scores for 3×2-meter remained high throughout the sequence, whereas syncopation scores for 2×3-meter monotonically increased along with the 2×2-meter used in the main analysis. This suggests that the sequences did not change between different meters, but gradually changed from providing clear cues to a 2×2 meter to containing little cues to any regular meter.

#### Cochlear model

Importantly, the main motivation for using the exact inversions of the regular-to-syncopated sequences to generate the syncopated-to-regular sequences was to ensure that the envelope spectra of the original and inverted sequence were identical (due to the properties of the Discrete Fourier Transform). This way, differences between the original and inverted sequences in the EEG response across corresponding segments can only be explained by recent stimulus history. To ensure that other nonlinearities in the auditory system (such as adaptation) were not likely to explain the differences between the original and inverted sequences in the EEG response, the stimuli were analysed with a cochlear model. The model consisted of a Patterson-Holdsworth ERB filter bank with 100 channels (Patterson and Holdsworth, 1996), followed by Meddis’ hair-cell model (Meddis, 1986), as implemented in the Auditory Toolbox for Matlab (Slaney, 1998). The output of this model is designed to approximate sound representation in the auditory nerve, after narrowband filtering at the level of cochlea and nonlinearities introduced at the hair-cell level (adaptation, compression). The output of the cochlear model for each trial and sequence direction was segmented into seven 14.4-sec-long segments with 50% overlap. To obtain exactly the same envelope signals across corresponding segments from the two sequence directions, segmenting started at event 13 for the regular-to-syncopated condition and at event 10 for the syncopated-to-regular condition (see Figure 1). The time-domain signals were averaged across trials separately for each 14.4-s segment and sequence direction, and transformed into the frequency-domain using fast Fourier transform (FFT, yielding a spectral resolution of 1/14.4 s, i.e. approximately 0.069 Hz). The resulting spectra were averaged across cochlear channels.

As depicted in Figure 3, none of the obtained spectra showed clear peaks emerging from the spectral background, except at the frequency of individual events (5 Hz), and half this rate (2.5 Hz). This was due to the fact that none of the patterns making up the sequences were repeated within the sequence, thus yielding no prominent periodicities in the sequences except those related to individual events and successions of two events. As the sequences gradually transformed from regular to syncopated, the prominence of the peak at 2.5 Hz decreased over the segments, and the spectral energy spread across other frequencies, thus indicating, as intended, the absence of prominent cues to any particular higher-order structure beyond the event rate.

**Figure 3.**
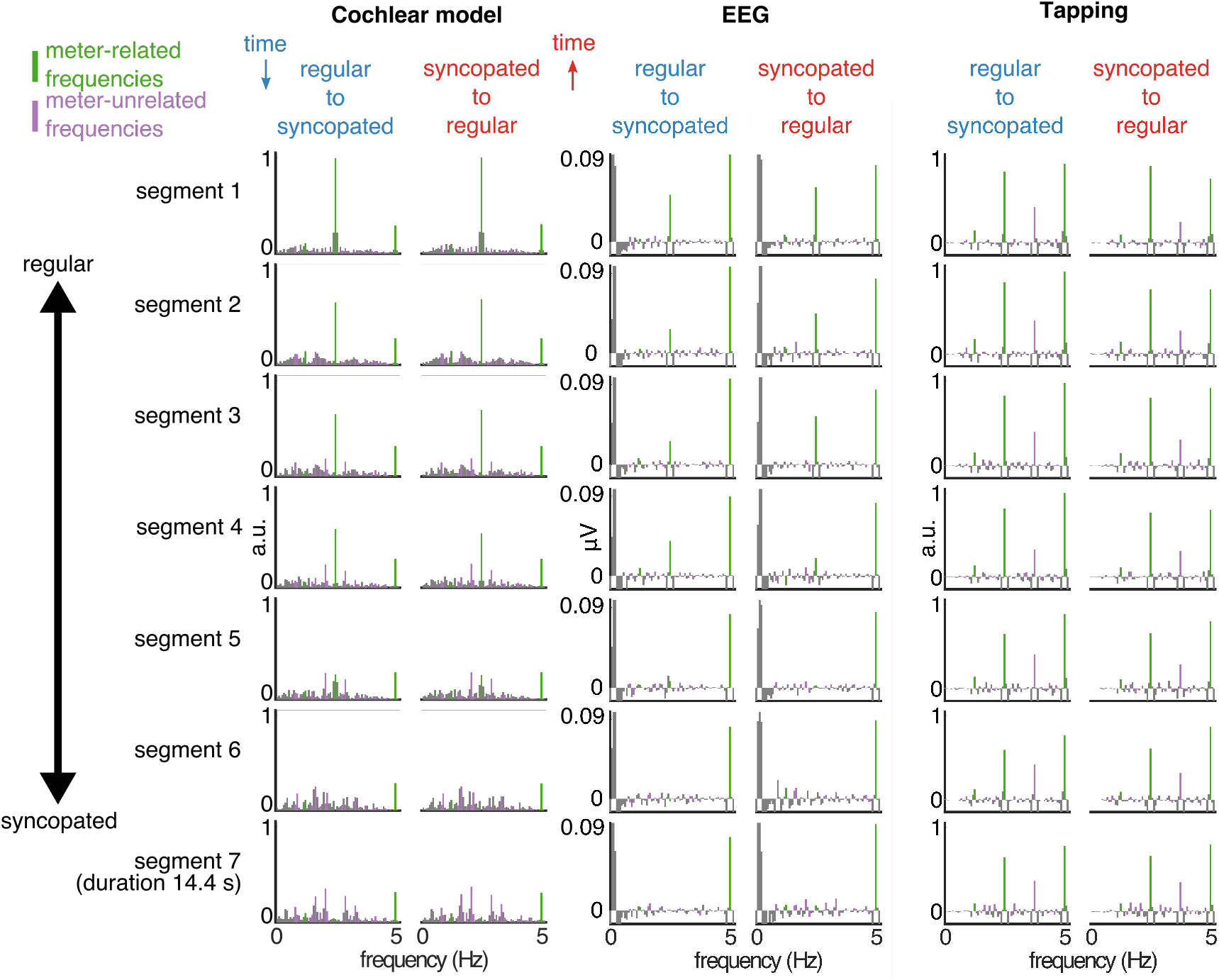
Cochlear model, EEG, and tapping spectra. The data are averaged across trials and plotted separately for each segment and sequence direction. The segments from the syncopated-to-regular condition are displayed in reverse order to facilitate comparison across conditions (this way the segments with the same stimulus envelope spectra are aligned). The cochlear model output (Left) shows highly similar spectra across sequence directions, with decreasing prominence of meter-related frequencies (green) and increasing prominence of meter-unrelated frequencies (purple) as the sequence changes from regular to syncopated. The EEG response (Middle) averaged across all channels and participants contains peaks at the frequencies present in the cochlear model output, with decreasing prominence of meter-related frequencies in the syncopated segments. The tapping response (Right) averaged across participants shows prominent peaks at meter-related frequencies even in the syncopated segments.

To make sure that the output of the cochlear model was not significantly different between sequence directions, especially at the frequencies related to the induced meter, we measured the amplitude at specific frequencies in the obtained spectra. These frequencies corresponded to different possible groupings of the events comprising the sequence, i.e., considering cycles of 12 events (0.416 Hz) and 16 events (0.312 Hz) and their harmonics up to 5 Hz (individual event frequency). From this set (N = 21 frequencies), a subset of frequencies was categorized as related to the induced meter (1.25, 2.5 and 5 Hz, as confirmed by the tapping session; see section Tapping analysis). These meter-related frequencies represent nested grouping of the individual event rate (5 Hz) by 2 (2.5 Hz) and 2 (1.25 Hz), thus corresponding to the meter used to construct the sequences (excluding the bar-level grouping, as for the syncopation score calculation). The rest of the frequencies were considered meter-unrelated. The amplitude at each frequency was extracted either at the exact frequency, if a bin was centred at that frequency (14 frequencies), or otherwise as a maximum value from the two closest bins. The 21 extracted amplitudes were z-scored as follows: (x − mean across the 21 frequencies)/SD across the 21 frequencies. This standardization evaluated the magnitude at each frequency relative to the other frequencies, and therefore allowed us to quantify how much a particular subset of frequencies (here meter-related frequencies) stood out relative to the whole set of frequencies. Because this measure is invariant to differences in unit and scale, it also enabled us to objectively measure the relative distance between stimulus representation at the earliest stages of the auditory pathway (estimated with the cochlear model) and the elicited EEG response.

The relative prominence of meter-related frequencies in the cochlear model output (considering the whole set of extracted frequencies) was calculated as a mean z score at 1.25, 2.5 and 5 Hz. These meter-related z scores were compared between the two sequence directions across segments to ensure that the inversion of the stimulus was unlikely to introduce significant differences in the prominence of meter frequencies at the earliest stages of the auditory pathway. For this comparison, the z-scored amplitudes were extracted in the way described above but separately for each trial (i.e. without first averaging across trials in the time domain), and fitted with a mixed model (fixed effects Direction and Segment). There were no significant differences between the original and inverted condition (main effect of Direction, F = 0.01, P = 0.92, BF_10_ = 0.15; interaction of Direction and Segment, F = 0.64, P = 0.7, BF_10_ = 0.07). This result suggests that nonlinearities at the early stages of the auditory pathway are unlikely to account for any effects of context in the EEG responses.

The same analyses performed on the 5 sequences used in the tapping session suggested similar differences in syncopation scores (higher syncopation score for regular-to-syncopated condition in segment 4, β = 4.6, t = 3.88, P = 0.002), and no significant effects involving the factor Direction for the analysis with cochlear model (Ps > 0.82, BFs_10_ < 0.25).

### Experimental design and procedure

The experiment consisted of an EEG and a tapping session directly following each other. In the EEG session, participants were presented with the 15 sequences and their inverted versions in random order with regular-to-syncopated and syncopated-to-regular trials alternating (counterbalanced across participants). Participants were seated in a comfortable chair with their head resting on a support, and asked to avoid any unnecessary movement. The support made contact with the head just below the most inferiorly positioned electrodes in order to prevent artifacts in the recorded EEG signals. Participants were asked to focus on the regular beat in the auditory stimuli, and after each trial, to rate (on a scale from 1 to 5) how difficult on average they thought it would be to tap along the beat in that trial. The ratings were taken as a subjective measure of difficulty in perceiving a clear meter in the sequences. Furthermore, to encourage attention to the temporal properties of the stimuli, participants were also asked to detect slight transient decrease of tempo randomly inserted in two additional trials that were not included in the analyses. Before the EEG session, the experimenter provided examples of beat in popular music and artificially constructed rhythms, to make sure participants understood the task.

After the EEG session, participants were presented with five additional sequences and the respective inverted versions, and were asked to tap the regular beat they perceived in the sequences using the index finger of the preferred hand. Tapping was performed on a custom-built response box containing a piezoelectric sensor that converted the mechanical vibrations of the box due to the impact of the finger into electrical signals, which were recorded as audio files.

### EEG recording and preprocessing

The EEG was recorded using a Biosemi Active-Two system (Biosemi, Amsterdam, Netherlands) with 64 Ag-AgCl electrodes placed on the scalp according to the international 10/20 system, and two additional electrodes attached to the mastoids. Head movements were monitored using an accelerometer with two axes (front-back and left-right) attached to the EEG cap and recorded as 2 additional channels. The signals were digitized at a 2048-Hz sampling rate and downsampled to 512 Hz offline.

The continuous EEG signals were high-pass filtered at 0.1 Hz (4th order Butterworth filter) to remove slow drifts from the signal. Independent component analysis (Bell and Sejnowski, 1995; Jung et al., 2000) was used to identify and remove artifacts related to eye blinks and horizontal eye movements based on visual inspection of their typical waveform shape and topographic distribution (2 components removed for 14 participants, 1 component for 18 participants). An independent component was only removed if it was within the first 10 components explaining the largest amount of variance. Channels containing excessive artifacts or noise were linearly interpolated using the 3 closest channels (1 channel interpolated for 2 participants, 4 channels for 1 participant). The cleaned EEG data were segmented into 57.6-s long epochs, starting from 2.4 s (event 13) for the regular-to-syncopated condition and from 1.8 s (event 10) for the syncopated-to-regular condition relative to the trial onset (due to padding with sound events, see above section Auditory stimulation and Figure 1). The data were then further segmented into seven 14.4-sec long segments with 50% overlap (as for the auditory stimulus analysis), re-referenced to the common average, and averaged across trials in the time domain separately for each sequence direction, segment, and participant. Time-domain averaging was performed to increase the signal-to-noise ratio of the neural response by cancelling signals that were not time-locked to the stimulus (Mouraux et al., 2011; Nozaradan et al., 2011, 2012). To assess the contribution of this processing step to our results, we also performed a control analysis without time-domain averaging, where the data were first transformed into the frequency domain (see section below) and the resulting spectra were averaged across trials. The EEG preprocessing was carried out using Letswave6 (www.letswave.org) and Matlab.

### Frequency-domain analysis of EEG response

For each participant, sequence direction, and segment, the EEG signals were transformed into the frequency domain using FFT. The obtained EEG spectra can be assumed to consist of a superposition of (i) responses to the stimulus concentrated into narrow peaks and (ii) residual background noise smoothly spread across the entire frequency range. To obtain valid estimates of the responses, the contribution of noise was minimized by subtracting, at each frequency bin, the average amplitude in the 2nd neighboring bin either side of it (Mouraux et al., 2011; Xu et al., 2017). A control analysis conducted on the EEG spectra obtained without noise subtraction yielded similar results to the analysis incorporating noise subtraction (see Results section), showing that this processing step alone could not explain our results. The noise-subtracted spectra were averaged across all channels to avoid electrode-selection bias and to account for individual differences in response topography.

To assess the relative prominence of the specific frequencies in the EEG responses elicited by the auditory stimuli, amplitudes at the 21 frequencies corresponding to different possible metric interpretations were then extracted from the spectra and z-scored in the same way as for the auditory stimulus analysis. The greater z score at a specific frequency indicates more prominent amplitude at that frequency relative to the whole set of 21 frequencies in the EEG response. Mean z-scored amplitude at frequencies related to the induced meter (5 Hz, 2.5 Hz and 1.25 Hz, as theoretically expected based on the sequence generation algorithm and as indicated by tapping analysis) was taken as a relative measure of selective synchronization of neural activity at the meter periodicities (control analysis with raw EEG amplitudes yielded similar results to the analysis with z scores, see Results section). The mean meter-related z-scored amplitudes were compared across sequence directions and segments, by fitting a mixed model (fixed effects Direction, Segment, and Musical Training). We expected to find a decrease in the prominence of meter-related frequencies in the segments with higher syncopation, as in the auditory stimulus. Importantly, we used additional post-hoc contrasts to test whether the EEG response was affected by the direction of the sequence by comparing the prominence of meter frequencies in segment one (most regular rhythm) to all subsequent segments, separately for each sequence direction. We hypothesized that in the regular-to-syncopated condition, the decrease would take place in segments with higher syncopation scores compared to the syncopated-to-regular condition. We also directly compared segments that had identical sound envelope spectra across sequence directions, to assess whether the EEG response at meter-related frequencies would be enhanced for particular segments in the regular-to-syncopated condition.

To further show that cochlear processing was unlikely to explain the effect of context in the EEG responses, the two signals were directly compared after standardization (z-scoring). In order to use the same processing pipeline for the EEG and cochlear model (see section Stimulus analysis), the cochlear model spectra were noise-subtracted (bin 2 on each side) before z-scoring the magnitudes across the meter-related and meter-unrelated frequencies. Subsequently, the difference in meter-related z scores between the cochlear model and the EEG response was calculated separately for each sequence direction, segment, and participant. The difference scores were compared between sequence directions, segments, and levels of musical training with a mixed model, and post-hoc contrasts compared the difference score between directions separately for each segment. Hence, if the EEG responses were fully explained by cochlear processing, the obtained scores should not significantly differ between the two sequence directions.

### Rating analysis

Ratings of subjective difficulty in perceiving a clear meter in the stimuli were averaged across trials, separately for each sequence direction and participant. To evaluate whether the subjective perception of meter was affected by sequence direction, the mean ratings were compared between the regular-to-syncopated and syncopated-to-regular condition by fitting a mixed model with Direction and Musical Training as fixed effects.

### Tapping analysis

Tap times were extracted by locating points in the continuous signal from the tapping sensor where the (i) amplitude was increasing, (ii) amplitude exceeded a threshold set manually for each participant, and (iii) the amount of time from the previous detected point was larger than a constant set manually for each participant. These points corresponded to the tap onsets, i.e. the times where the finger hit the response box. To quantitatively evaluate the meter periodicities participants synchronized to, the median inter-tap interval (ITI) was calculated separately for each sequence direction and participant. The value was then compared to three possible meters each consisting of 3 nested periodicities (nested grouping by 2×2; 2×3; and 3×2 events, corresponding to periods 200, 400, 800 ms; 200, 400, 1200 ms; and 200, 600, 1200 ms, respectively) by taking the minimum percent difference between the median ITI and the three possible periodicities comprising each meter. This minimum difference score was compared across meters and sequence directions using a mixed model. The meter that yielded the smallest difference score was considered to be the meter predominantly induced by the stimulus construction method.

Furthermore, similar to the auditory stimuli and EEG responses, tapping responses were also analyzed in the frequency domain to evaluate synchronization at meter-related frequencies at the level of behavioral output. Continuous signals from the response box recorded during the tapping session were segmented the same way as the EEG signals and transformed into the frequency-domain separately for each trial. The contribution of background noise was minimized, as for the EEG, by subtracting the average magnitude in the 2nd neighboring bin either side of each frequency-bin. The resulting spectra were averaged across trials, and magnitudes at meter-related and meter-unrelated frequencies were extracted and z-scored as for the EEG analysis. Mean z-scored amplitudes at meter-related frequencies were compared across segments, sequence directions, and levels of musical training, by fitting a mixed model. The persistence of the tapping synchronization across different amounts of syncopation was assessed using post-hoc contrasts that compared the prominence of meter-related frequencies in the first segment to all subsequent segments. To further understand the evolution of the tapping response over segments, the prominence of meter frequencies was also compared across all pairs of successive segments.

### Head movement analysis

To evaluate the extent to which unintentional head movement artifacts could explain the observed EEG results, the data from the accelerometer were segmented the same way as EEG signals and transformed into the frequency-domain separately for each axis. The resulting spectra were averaged across the two axes, and mean magnitudes at meter-related frequencies were extracted and further analyzed as for the EEG responses.

### Statistical analyses

The statistical analyses were performed using linear mixed models with lme4 package in R (Bates et al., 2014). Each participant was included as a random-effect intercept (in case of stimulus analyses, intercept was modeled as a random variable across trials). For models including the factor Segment as a fixed effect, the order of segments from the syncopated-to-regular condition was always reversed in order to compare responses to the inverted version of the same acoustic stimulus. Post-hoc multiple comparisons were computed using emmeans package (Lenth, 2018). The Kenward-Roger approach was used to approximate degrees of freedom and Bonferroni correction was used to adjust for multiple comparisons. Complementary to the null-hypothesis significance tests with mixed models, we also calculated Bayes factors to quantify the evidence in favor of the alternative hypothesis over the null hypothesis (BF_10_), as implemented in the package BayesFactor for R (Morey and Rouder, 2014).

## Results

### Behavioral results

#### Ratings

The ratings of subjective difficulty at perceiving a clear meter did not differ significantly across sequence directions (F = 0.10, P = 0.75, BF_10_ = 0.26). There was also a trend towards the main effect of musical training (F = 3.06, P = 0.09, BF_10_ = 0.86), suggesting that some musicians tended to find it easier to perceive a clear beat in the sequences. However, because the ratings were given for the whole trials, this measure might be too coarse to capture possible local and transient effects of context.

#### Tapping

The tapping task confirmed theoretical expectations about the meter periodicities induced by the auditory stimulus sequences. The difference between the median ITI and possible meter periodicities varied significantly across the different possible meters (F = 19.65, P < 0.0001, BF_10_ > 100). Post-hoc comparisons showed that the median ITI was significantly closer to the 2×2-meter than the 2×3-meter (β = −13.22, t = −5.39, P < 0.0001) and 3×2-meter (β = −13.54, t = −5.52, P < 0.0001). These results further justify the selection of meter-related frequencies (5Hz, 5 Hz/2 and 5 Hz/4, corresponding to the individual event rate, grouping by two events, and grouping by four events, respectively) for the frequency-domain analyses.

The spectra of continuous signals from the tapping sensor exhibited prominent peaks at meter-related frequencies (Figure 3). As depicted in Figure 4, the prominence of these frequencies in the tapping spectra evolved across segments differently for musicians and non-musicians (F = 8.61, P < 0.0001, BF_10_ > 100). When comparing the two groups separately for each segment, meter frequencies were more prominent for musicians in segments 6 (β = 0.82, t = 4.54, P = 0.0002) and 7 (β = 1, t = 5.59, P < 0.0001). This was due to the fact that for non-musicians, meter frequencies significantly decreased in segments 5 (β = −0.35, t = −3.35, P = 0.01), 6 (β = −0.84, t = −8.10, P < 0.0001) and 7 (β = −1.08, t = −10.38, P < 0.0001) when compared to segment 1, while for musicians there was only a trend towards a decrease in the last (i.e. most syncopated) segment (β = −0.30, t = −2.89, P = 0.05). Another set of contrasts sequentially comparing pairs of successive segments revealed a sharp decrease between segment 5 and 6 (β = −0.49, t = −4.74, P < 0.0001) for non-musicians, but not for musicians (Ps = 1.00). This indicates that the ability of non-musicians to synchronize their tapping at meter frequencies deteriorated significantly once the syncopation level exceeded a critical level. Together, these results suggest that even though musicians were influenced by syncopation in the stimulus, their ability to tap along with the regular meter decreased more steadily as syncopation increased.

**Figure 4.**
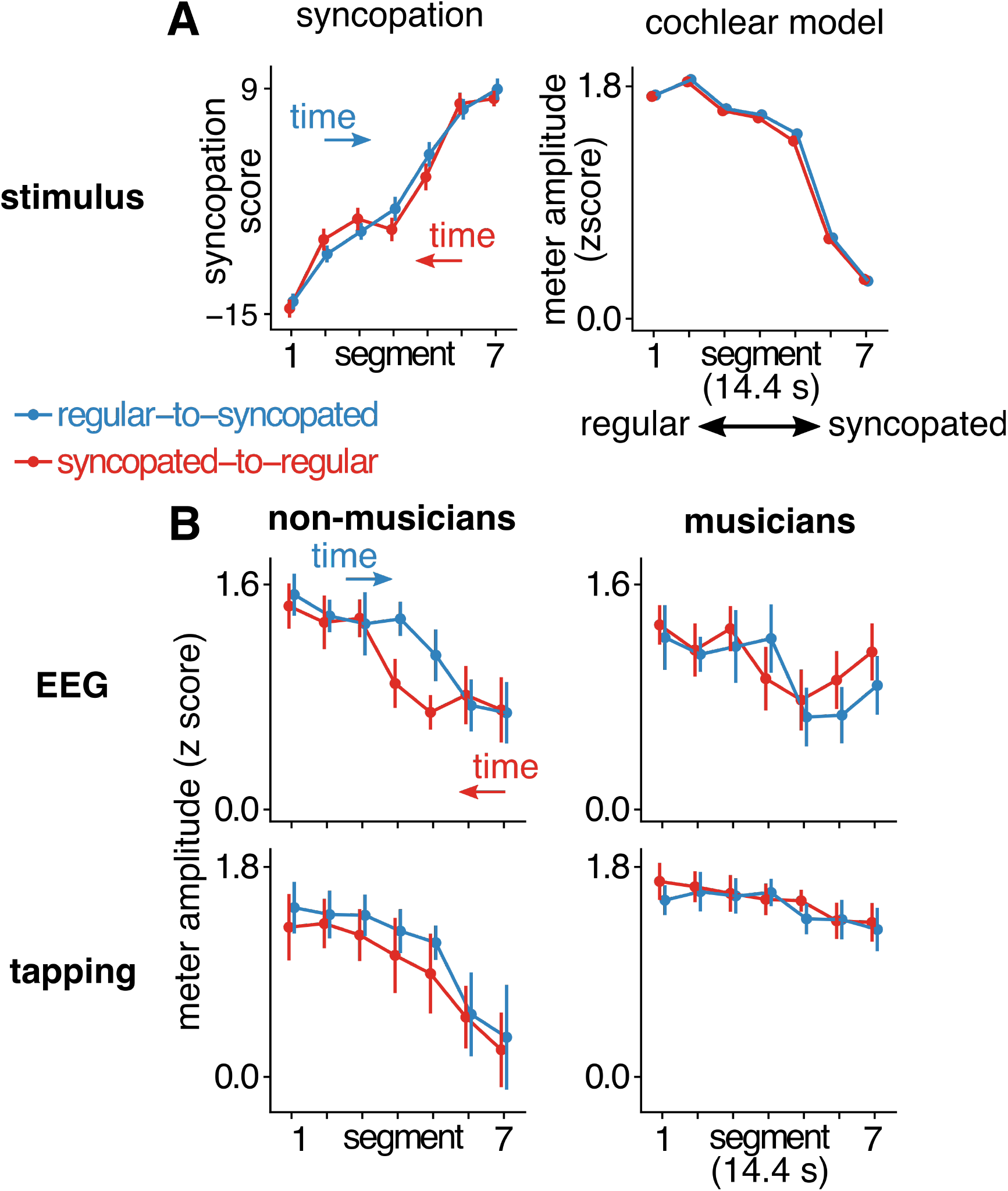
Analysis of stimulus, EEG, and tapping response. The order of segments in the syncopated-to-regular condition (red) is reversed to aid the comparison of segments with identical stimulus envelope spectra across conditions. Mean values are shown as points, and error bars represent 95% confidence interval (Morey, 2008). (A) Analysis of the stimulus. (Left) Syncopation scores averaged across trials separately for each condition, with arrows indicating the direction of time for each condition. (Right) Mean z-scored amplitudes at meter-related frequencies in the cochlear model output. As intended, the prominence of meter frequencies decreased as the syncopation of the sequence increased. (B) Mean z-scored amplitudes at meter-related frequencies for the EEG responses (Top) and tapping responses (Bottom), plotted separately for non-musicians (Left) and musicians (Right). Non-musicians showed enhanced EEG responses at meter frequencies in the middle segments of the regular-to-syncopated condition (blue), and the prominence of meter frequencies in their tapping decreased rapidly with increasing syncopation. The EEG responses of musicians were similar across conditions and their tapping showed prominent meter frequencies even in the syncopated segments.

There was also a significant interaction between musical training and condition (F = 6.30, P = 0.01, BF_10_ = 2.16). While the overall prominence of meter frequencies was larger in the tapping of musicians for both sequence directions, this difference was more pronounced in the syncopated-to-regular condition (β = 0.58, t = 3.70, P = 0.001) than the regular-to-syncopated condition (β = 0.39, t = 2.45, P = 0.04). This was due to the fact that non-musicians showed overall smaller prominence of meter frequencies in the syncopated-to-regular condition (β = 0.15, t = 2.61, P = 0.02). This result provides some evidence for improved tapping to the meter when the rhythm evolved from regular to syncopated. However, since the Bayes factor is smaller than 3, caution is called for when drawing conclusions from this result (Lee and Wagenmakers, 2014).

### Frequency-domain analysis of EEG

EEG responses were elicited at frequencies that were expected on the basis of the auditory stimulus analysis (Figure 3), with typical fronto-central topographies (Figure 5), as previously observed for responses to repeating auditory rhythms (Nozaradan et al., 2012; Lenc et al., 2018).

**Figure 5.**
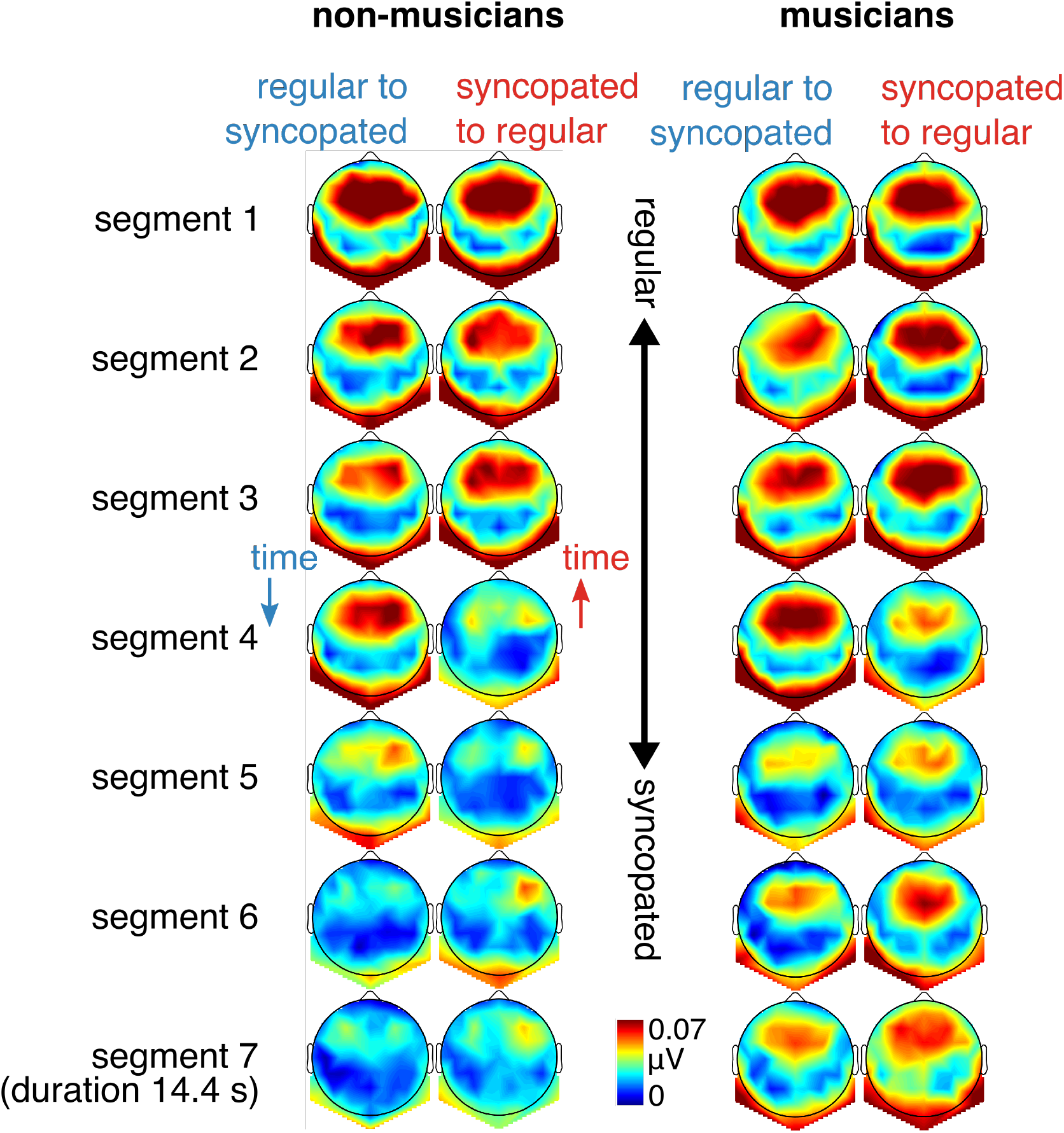
Topographies of the mean EEG amplitude at meter-related frequencies. Scalp distributions of responses across conditions and segments are shown separately for non-musicians (Left) and musicians (Right).

The main aim of the current study was to examine the effect of context on the relative amplitude of EEG responses at meter-related frequencies (Figure 4). The direction of the sequence affected the prominence of meter-related frequencies (mean z-scored amplitudes) in the EEG response (interaction between Direction and Segment, F = 4.26, P = 0.0004, BF_10_ = 33.70). Directly contrasting the corresponding segments between the two sequence directions revealed significantly larger meter frequencies for segment 4 (β = 0.37, t = 4.16, P = 0.0002) in the regular-to-syncopated condition compared to the opposite sequence direction. This was due to greater persistence of the response in the regular-to-syncopated condition, as syncopation increased. Table 1 shows the response across segments compared to the first segment, separately for musicians and non-musicians. For non-musicians, the response significantly decreased in segment 5, 6, and 7 in the regular-to-syncopated condition. However, for the syncopated-to-regular condition, there was a significant decrease already in segment 4, followed by segment 5, 6, and 7. Similar pattern of results can be observed for musicians (significant decrease in segments 5 and 6 for regular-to-syncopated and segments 4, 5, and 6 in the opposite direction). In other words, in the segment with medium amount of syncopation, the meter-related frequencies were more prominent in the EEG when regular, as opposed to syncopated, input preceded this segment. Additionally, even though the three-way interaction between sequence direction, segment, and musical training was not significant (F = 0.71, P = 0.64, BF_10_ = 0.07), the context effect seemed to be less pronounced in the group of musicians.

**Table 1.**
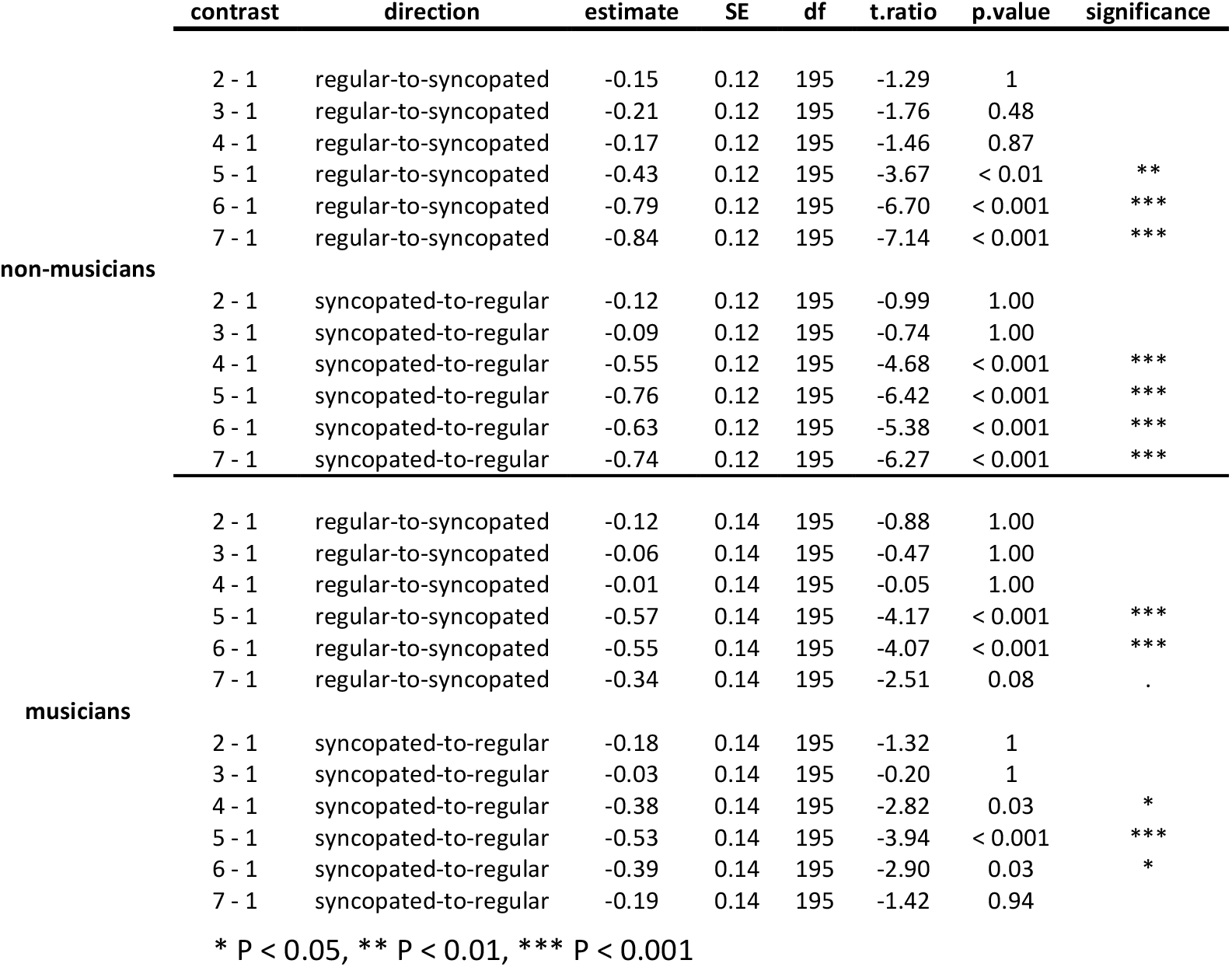
Prominence of meter-related frequencies in the EEG response compared between the first and all subsequent segments.

Furthermore, there was an interaction between musical training and segment (F = 4.35, P = 0.0003, BF_10_ = 41.70). However, this effect seemed primarily driven by greater selective response at meter-related frequencies in segment 7 for musicians, which did not reach significance in the post hoc contrasts (β = 0.30, t = 2.57, P = 0.08). Finally, musical training interacted with sequence direction (F = 9.03, P = 0.003, BF_10_ = 6.51). However, post hoc contrasts did not reveal significant differences between musicians and non-musicians in either condition (Ps > 0.13).

Direct comparison between the cochlear model and EEG indicated that the results are unlikely to be fully explained by nonlinearities in the early stages of the auditory pathway. The difference in prominence of meter-related frequencies between EEG and cochlear model significantly depended on the direction of the sequence (interaction between Direction and Segment, F = 8.42, P < 0.0001, BF_10_ > 100) for segment 4 (β = 0.36, t = 5.91, P < 0.0001).

Similar to the main analysis, there was an interaction of musical training and segment (F = 9.34, P < 0.0001, BF_10_ > 100), driven by greater response and meter frequencies for musicians in segment 7 (β = 0.30, t = 2.99, P = 0.03). Additionally, there was an interaction between musical training and sequence direction (F = 19.4, P < 0.0001, BF_10_ > 100). Again, directly contrasting musicians and non-musicians for each condition did not yield significant differences (Ps > 0.12).

Together, these results indicate that even after accounting for the response variability explained by the cochlear model, EEG response at meter-related frequencies was significantly affected by sequence direction.

### Control analyses of EEG

#### Raw amplitudes

Z-scoring EEG amplitude values across a set of frequencies was used to assess the selective variations in the amplitudes at meter-related frequencies in a manner that minimized the contribution of the overall gain. However, to demonstrate that this standardization procedure was not responsible for the context effect observed in the current study, we carried out a control analysis of the EEG response without any normalization, i.e. using raw amplitude values from the EEG spectra averaged over meter-related frequencies as a dependent measure. The amplitude at meter frequencies was affected by sequence direction (interaction between Direction and Segment, F = 6.97, P < 0001, BF_10_ > 100), and this was due to significantly larger amplitude at meter frequencies in the regular-to-syncopated condition for segment 4 (β = 0.37, t = 4.16, P = 0.0003). There was also a significant interaction between musical training and segment (F = 4.44, P = 0.0002, BF_10_ = 39.56), due to the marginally higher prominence of meter frequencies in segment 7 for musicians (β = 0.30, t = 2.57, P = 0.08). These results suggest that z-scoring EEG amplitudes alone is not likely to explain the current results.

#### Without noise subtraction

The prominence of meter-related frequencies extracted from EEG spectra without noise subtraction was affected by sequence direction (interaction between Direction and Segment, F = 5.30, P < 0.0001, BF_10_ > 100), due to greater meter z score in segment 4 (β = 0.22, t = 3.30, P = 0.007). Interaction between musical training and segment also reached significance (F = 2.72, P = 0.01, BF_10_ = 1.45), but there was no significant difference for either segment separately (Ps > 0.47). These results suggest that noise subtraction alone is not likely to explain the current results.

#### Without time-domain averaging

The control analysis using z-scored amplitudes extracted from the EEG spectra obtained without first averaging across trials in the time domain yielded results that were different from the main analysis (which included time-domain averaging). Even though there was a trend towards an interaction between sequence direction and segment (F = 2.03, P = 0.06, BF_10_ = 0.38), post hoc contrasts did not reveal differences between conditions in any segment (except for a trend towards larger response in syncopated-to-regular condition for segment 7; β = −0.28, t = −2.54, P = 0.08). There was also a significant interaction between musical training and segment (F = 2.58, P = 0.02, BF_10_ = 1.20), but post hoc contrasts did not reach significance (Ps > 0.21). Considering the low Bayes factors, these results do not provide unequivocal evidence supporting the reliability of the context effect we observed in the main analysis. This lack of effect could be related to the low signal-to-noise ratio of the EEG responses obtained without time-domain averaging across trials. To assess this possibility, we computed the ratio between the amplitude at each meter-related frequency and its 2nd neighboring bins, taken from EEG spectra without noise subtraction obtained either with or without time-domain averaging. The ratios averaged across meter frequencies, sequence directions, and segments, were significantly larger when time-domain averaging was included (paired t-test, t = 15.16, P < 0.0001, effect size: Cohen’s d = 2.68). This result underscores the relevance of time-domain averaging to obtain robust estimates of time-locked responses from EEG signals, particularly when these responses have small amplitudes (see Figure 3).

### Head movement analysis

The prominence of meter-related frequencies in the head movement data did not significantly differ between sequence directions (main effect of Direction, F = 0.82, P = 0.37, BF_10_ = 0.16). There was a significant interaction between Direction and Segment (F = 2.50, P = 0.02, BF_10_ = 3.89). However, testing individual segments for differences between regular-to-syncopated and syncopated-to-regular condition did not yield any significant contrasts (Ps > 0.27). Together, this control analysis suggests that the observed EEG effects are unlikely to be explained by head movement artifacts.

## Discussion

Our results show direct evidence that the neural tracking of rhythmic input is affected by recent auditory context. Specifically, we observed in the EEG a selective enhancement of meter-related frequencies that persisted even when the periodic acoustic cues guiding meter perception were gradually degraded in the stimulus. Conversely, these meter-relevant frequencies were less prominent in the neural response when the preceding acoustic input lacked periodic cues to guide meter perception. Moreover, this context effect was more prominent in the group of participants with no formal musical training, who also demonstrated lower ability to tap along with a stable meter when it was not supported by periodic temporal cues in the stimulus. In contrast, the context effect was weaker in the group of musicians whose ability to maintain a meter was overall much more robust to stimulus ambiguities, as observed in the tapping session. Together, these results show that neural selection of functionally-relevant features is continuously shaped by incoming and recent sensory input as well as preexisting perceptual templates.

### Dissociation between selective enhancement of meter frequencies and acoustic input: evidence from syncopated rhythm

Our results provide evidence supporting the view that neural responses to rhythmic sound sequences do not stand in a one-to-one relationship with the rhythmic modulations of the acoustic stimulus. Rather, these neural responses exhibit a degree of dissociation from these acoustic features, which could support the emergence of perceptual representations that are invariant with respect to the input (Ding and Simon, 2012; Zion Golumbic et al., 2013; Nozaradan et al., 2017a; Kuchibhotla and Bathellier, 2018). In other words, different rhythmic patterns can give rise to the perception of the same pulse-like meter, and the selection and maintenance of these perceived periodicities across a wide range of inputs rely on a flexible combination of sensory and endogenous signals, leading to prominent neural responses at meter frequencies regardless of their prominence in the input.

Our results also add to the evidence that these neural processes depend on prior experience at different time scales (Chemin et al., 2014; Cirelli et al., 2016; Stupacher et al., 2017). Previous work has shown that sensory-motor synchronization emphasizing a certain metric interpretation of an ambiguous rhythmic input can significantly shape neural processing during subsequent listening of the same input, in the form of enhanced neural responses at the frequencies corresponding to the meter the listener previously moved to (Chemin et al., 2014). Here we move a step forward by showing that even without movement, previous exposure to an auditory rhythmic input containing prominent meter periodicities can lead to persistent neural emphasis on these meter frequencies even if they are degraded in the subsequent input.

This demonstrates hysteresis in neural synchronization to the periodic meter, i.e. dependence of the response on recent past (Large, 2000; Kleinschmidt et al., 2002; Melloni et al., 2011; Bouvet et al., 2019). Such nature of the response might point towards similarities between musical meter processing and other high level perceptual phenomena such as speech or visual object recognition, where prior exposure to the specific stimulus can facilitate perceptual disambiguation of that stimulus in impoverished conditions (Dolan et al., 1997; Davis et al., 2005; Holdgraf et al., 2016). Accordingly, the prolonged perception of picture identity as it is slowly going out of focus (Bruner and Potter, 1964) could be equivalent to the persistence of meter perception as a rhythm gradually becomes more and more ambiguous. Such similarities would be in line with the predictive nature of perceptual processing and the importance of endogenous information (such as prior knowledge and expectations) in shaping the processing of sensory signals across domains (de Lange et al., 2018; Koelsch et al., 2019).

### Short-lived time constant for context effect on meter perception

In the current study, the contextual enhancement of meter-related frequencies was relatively short-lived, i.e. lasting ~14 seconds. Moreover, the effect did not carry over across trials. These observations demonstrate that the influence of prior acoustic context on EEG responses might have a short time constant, only affecting the processing of directly following rhythmic material. Such short-lived integrative mechanism would thus make the system both robust to momentary changes in the rhythmic structure (e.g. syncopation) and flexible enough to adjust meter perception under persisting counterevidence from the sensory input (London, 2004).

This short time constant observed here could also be due to the sequence design combined with a context effect restricted to inputs up to a certain level of meter ambiguity. Indeed, in the current study, the sequences were designed to provide a systematic increase of syncopation all the way to complete meter ambiguity. However, the limited ability to perceive and synchronize to a stable meter when listening to highly syncopated rhythms (Fitch and Rosenfeld, 2007; Vuust et al., 2018) suggests a restricted capacity of the system to endogenously generate a set of metric periodicities (Nozaradan et al., 2012), especially in cases of strong mismatch between the stimulus acoustic structure and perceptual metric organization. Consequently, this would confine the effects of prior context to inputs with medium amounts of syncopation, thus explaining why we did not observe selective enhancement of meter frequencies in response to the most syncopated sections of the sequences.

The relatively short time constant observed here could also be due to context effect restricted to inputs up to a certain degree of similarity with the recently heard material, in line with research on serial effects in vision (Manassi et al., 2017; Cicchini et al., 2018). Such a mechanism would bias the neural response towards recent past in case of small difference between the current and directly preceding input. This bias would be absent in case of abrupt changes in the input, supporting adaptive changes of perceptual organization in light of substantial sensory evidence.

Our results thus pave the way for future investigation of these neural biases, and whether they can be transferred between spectral channels and perceptual objects, to further elucidate how the brain achieves stable meter perception from acoustically complex sequences such as naturalistic music.

### Underlying neuronal properties and neural network

The context effect observed here could be supported by an increased sensitivity of neurons to modulation frequencies that were prominent in the recent input. A similar mechanism has been proposed in the visual system, where responses to a stimulus can be biased towards recently seen stimuli already in primary visual areas (St. John-Saaltink et al., 2016). In the auditory system, there is mounting evidence for input-output properties of neurons showing sensitivity to recently heard stimuli across the auditory pathway, in particular in auditory cortices (Shamma and Fritz, 2014; Holdgraf et al., 2016; David, 2018). These mechanisms are thus compatible with the effects of context observed here, biasing the input-output transformation of rhythm in favor of meter frequencies.

These contextual changes in input-output properties of auditory neurons can also be modulated by higher-level associative and/or motor areas (Patel and Iversen, 2014; Nozaradan et al., 2017b). Such mechanisms have been observed in invasive animal studies where frontal regions have been shown to modify the encoding of behaviorally-relevant sensory inputs in the primary auditory cortex (Fritz et al., 2010; Winkowski et al., 2016). Relatedly, responses to rhythmic inputs in auditory cortices might be modulated through a number of bidirectional functional connections with subcortical and cortical structures, such as basal ganglia, premotor cortex and supplementary motor area, that have been implicated in meter processing (Patel and Iversen, 2014; Merchant et al., 2015). For example, there is evidence that the basal ganglia are involved in maintaining the metric organization established by directly preceding context (Grahn and Rowe, 2013), and overall in predictive timing (Grahn and Rowe, 2009; Teki et al., 2012; Merchant et al., 2015). This structure also contributes to the selective amplification of meter frequencies when listening to syncopated rhythms (Nozaradan et al., 2017b). Therefore, basal ganglia activity could play a role in biasing neural responses to incoming rhythmic inputs in favor of a recently induced meter, in line with the context effects observed here.

Another mechanism that potentially contributed to effects of recent context in the present study could be a top-down modulation of stimulus processing in auditory cortices exerted by frontal motor-planning areas (Patel and Iversen, 2014; Rimmele et al., 2018). Recently, it has been shown that frontal regions causally modulate speech envelope tracking in auditory cortices through low-frequency neural oscillations, implementing dynamically updated predictions that enhance encoding of slow speech modulations in primary sensory areas (Park et al., 2015, 2018). A context-dependent, predictive enhancement of meter-related frequencies when listening to musical rhythm could be therefore implemented in a similar vein, with frontal areas (i) modulating the processing of rhythmic input in auditory regions, and/or (ii) directly contributing to the scalp-recorded EEG signal as measured in the present study.

### Influence of preexisting perceptual templates in musicians

Recent context significantly shaped neural responses to incoming input in the group of musically naïve participants, but this effect seemed weaker in participants with musical training. This result could be related to the robustness of preexisting internal templates of meter in musicians, as also suggested in their ability to maintain stable tapping to the meter even in the most syncopated parts of the sequences. A trend towards similar effect was also observed in the EEG responses, whereby meter frequencies tended to be larger for musicians in the most complex segment irrespective of sequence direction. Musical training generally leads to superior precision of meter representation (Rüsseler et al., 2002; Brochard et al., 2003; Geiser et al., 2010; Lappe et al., 2011), with a high degree of invariance with respect to the rhythmic stimulus (Repp, 2007; Repp et al., 2008). Such robust ability to internally generate metric templates (Grahn and Rowe, 2009) could make musicians less influenced by incoming and recent input compared to nonmusicians. However, the effect of recent context was not completely absent in our heterogeneous group of musicians, thus calling for future investigations of how musical training in its different forms (Vuust et al., 2012; Bianco et al., 2018) affects meter perception and synchronization, and the neural processing of complex rhythms.

### Context effects in tapping

In contrast to the neural response, we did not observe an effect of recent context in sensory-motor synchronization during the tapping session. While the stimulus sequences used for the tapping task were not exactly identical to the sequences used for the EEG session, they were constructed with the same algorithm, and thus could be expected to elicit similar neural processes that would subsequently modulate overt behavior.

However, synchronized movement can directly (Nozaradan et al., 2013, 2016b; Morillon and Baillet, 2017; Yon et al., 2018) and prospectively (Lahav et al., 2007; Chemin et al., 2014) affect sound processing in the brain. Indeed, there is an increasing amount of evidence suggesting a bidirectional interaction whereby not only perception drives action, but overt action can affect perception (Maes et al., 2014; Witt, 2018). Accordingly, synchronized tapping has been shown to sharpen time prediction (Manning et al., 2016) and facilitate extraction of a periodic pulse-like beat from complex rhythmic sequences similar to the stimuli used in the current study (Su and Pöppel, 2012). Overt movement could therefore help selecting and maintaining an endogenous meter when the stimulus becomes ambiguous, thus attenuating the influence of recent auditory context. Importantly, these observations suggest that even though finger tapping might be useful to assess *what* pulse frequency people perceive (e.g. Nozaradan et al., 2012), in certain cases there may not be a one-to-one relationship between perception with and without movement.

Alternatively, the lack of context effects in tapping might be due to the smaller number of trials in the tapping session compared to the EEG session. The marginally significant main effect of sequence direction observed in nonmusicians’ tapping suggests that the location of the prior-context advantage within the sequences could have been variable across trials, yielding generally increased performance in the regular-to-syncopated sequence after averaging. These possibilities remain to be investigated with larger samples allowing for more detailed tapping analyses.

### Context effects and phase consistency

Another noteworthy result was the absence of context effects when the EEG responses from individual trials were directly transformed into the frequency-domain instead of first averaging in the time-domain. Averaging in the time-domain has been used across a number of EEG studies, as it increases signal-to-noise ratio of the responses to the input by cancelling out activity that is not time-locked to the input, and thus not aligned across trials (Picton, 2010; Mouraux et al., 2011; Nozaradan et al., 2011, 2018; Luck, 2014). An important assumption of this processing step is that the phase of the neural response of interest is constant across trials. If this is not the case, time-domain averaging can have detrimental effects on the amplitudes of individual frequency components in the average waveform (Lyons, 2011).

The extent to which the phase of neural response elicited by a rhythmic stimulus depends on the phase of the perceptual metric organization (i.e. the alignment of the metric pulses with respect to the input) remains unknown. While such link has been previously observed for transient EEG responses and modulation of high-frequency oscillatory activity (Iversen et al., 2009; Schaefer et al., 2011), the frequency-tagging approach might capture distinct dynamics of these neural responses, thus limiting direct comparisons across these methods. However, if there was a systematic relationship between perceptual and neural phase, this could offer an alternative explanation to the results of the current study. Specifically, the phase of the neural response could be more consistent across trials in the regular-to-syncopated sequences where the acoustic structure at the beginning of the sequences provided clear anchor points to the perceived meter. In the syncopated-to-regular sequences, in contrast, the beginning portions of the sequences did not favor particular meter frequencies or phases, making the internally generated meter more variable across trials. This differential consistency of meter phases across trials between the two sequence directions could thus account in part for the significant difference in EEG amplitude at meter frequencies observed between the sequence directions after time averaging.

### Conclusion

Our results provide direct evidence about how temporal context established though recently encountered auditory rhythmic stimuli can substantially modulate neural processing at subsequent time points. In particular, these results demonstrate that selective neural synchronization to metric periodicities persists over time when induced by a recent stimulus. We propose that this hysteresis in neural response to musical rhythm could represent a neurobiological basis for the remarkable flexibility and stability of meter perception relative to the acoustic input, and for the way music is often composed and performed. These observations also demonstrate how perception involves endogenous processes of short- and long-term neural plasticity continuously updating selection of functionally-relevant features.

